# Regulation of Perisomatic Synapses from Cholecystokinin Basket Interneurons through NrCAM and Ankyrin B

**DOI:** 10.1101/2024.11.04.621872

**Authors:** Erik N. Oldre, Barrett D. Webb, Justin E. Sperringer, Patricia F. Maness

## Abstract

The perisomatic region of cortical pyramidal neurons (PNs) integrates local and long-range inputs and regulates firing. This domain receives GABAergic inputs from cholecystokinin (CCK)- and Parvalbumin (PV)-expressing basket cells (BCs) but how synaptic contacts are established is unclear. Neuron-glial related cell adhesion molecule (NrCAM) is a homophilic transmembrane protein that binds the scaffold protein Ankyrin B. Here we show that NrCAM and Ankyrin B mediate perisomatic synaptic contact between CCK-BCs and PNs in mouse medial prefrontal cortex (mPFC). Immunolabeling of CCK-BC terminals for vesicular glutamate transporter-3 (VGLUT3) or vesicular GABA transporter (VGAT) revealed a significant decrease in CCK-BC synaptic puncta on PN soma in NrCAM-null mice, however no decrease in PV-BC puncta or cell loss. VGLUT3+ CCK-BC puncta were also decreased by Ankyrin B deletion from PNs in Nex1Cre-ERT2:Ank2^flox/flox^:EGFP mice. A novel CCK-BC reporter mouse expressing tdTomato (tdT) at the Synuclein-γ (*Sncg*) locus showed NrCAM localized to Sncg+ CCK-BCs, and to postsynaptic PN soma in Nex1Cre-ERT2:Ank2^+/+^:EGFP mice. Results suggest that NrCAM and Ankyrin B contribute to the establishment of connectivity between CCK-BCs and excitatory neurons of the mPFC.

## 1. Introduction

Establishment of a proper balance of excitatory and inhibitory (E/I) connectivity is achieved during synaptic development of cortical networks, and perturbation of this balance is implicated in neurodevelopmental disorders such as autism spectrum disorder (ASD) [1]. Pyramidal neurons (PNs) are the principal excitatory cells of the mammalian cerebral cortex. PNs transform synaptic inputs into action potentials that stimulate long- and short-range targets. The firing rates of PNs are tuned by GABAergic interneuron subtypes that form synapses at different subcellular domains. The perisomatic region of cortical PNs (soma, proximal dendrite, axon initial segment) is the key compartment for integrating diverse inputs and producing action potentials. GABAergic interneurons that target this domain regulate firing and synchronize a large array of PNs. The perisomatic domain of PNs receives almost exclusively GABAergic inputs derived from cholecystokinin (CCK)- and Parvalbumin (PV)-expressing basket cells (BCs) [2, 3]. PV-BCs are fast spiking interneurons that regulate timing and oscillations important for PN synchrony, whereas CCK-BCs are regular spiking interneurons that restrain firing, thus impacting mood, emotion, and working memory retrieval [4–7]. CCK-BCs expressing vesicular glutamate transporter 3 (VGLUT3) are unique in using both inhibitory and excitatory transmitters, GABA and glutamate, for functional neurotransmission onto postsynaptic PNs [8]. Normally, GABAergic inhibition is predominant, but glutamatergic transmission is activated under conditions of GABAergic restriction [8]. CCK-BCs are particularly important in the prefrontal cortex (PFC) [9, 7], which has essential roles in integrating inputs within circuits that regulate cognitive and emotional behaviors [10–14]. Although much is known about PV- BCs [5], CCK-BC development and function are less well understood, in part due to lack of appropriate transgenic lines.

Neuron-glial related cell adhesion molecule (NrCAM) is a transmembrane glycoprotein of the L1 family, whose members mediate diverse aspects of neural development, including cell adhesion, neuronal process growth, dendritic spine remodeling, and synaptogenesis [15]. The extracellular region of NrCAM has six Ig-like domains and five fibronectin III (FN) repeats, which engage in *trans* homophilic and heterophilic adhesion between cells. NrCAM mediates pruning of excess dendritic spines and excitatory synapses that are initially overproduced during postnatal development [16, 17]. Accordingly, NrCAM-null mice exhibit elevated levels of excitatory synapses and increased frequency of excitatory neurotransmission in neocortical PNs [18]. However, inhibitory properties have not been fully investigated in these mice. The conserved cytoplasmic domain of NrCAM reversibly engages Ankyrin B (AnkB), an intracellular spectrin-actin scaffold protein encoded by the high confidence ASD risk gene *Ank2* (SFARI Gene Database). AnkB also localizes the voltage-gated sodium channel Nav1.2 to the PN dendritic membrane important for back-propagation of action potentials [19]. Genomic studies revealed NrCAM variants associated with autism [20], and behavioral testing in NrCAM-null mice has demonstrated a reduction in learning and social behaviors relevant to autism [21].

An *in vivo* BioID-MS approach identified neuronal NrCAM in a proteome of inhibitory interneuron synapses from mouse brain using the GABAergic synaptic organizer, Gephyrin, as a bait protein [22]. A similar approach netted NrCAM in a proteome enriched for neuron-astrocyte junctions [23]. Depletion of NrCAM from astrocytes or neurons by CRISPR/Cas9 editing decreased inhibitory synapse formation in mouse visual cortex [23]. Astrocytic NrCAM deletion increased the rise time of miniature inhibitory postsynaptic potentials (mIPSCs) suggesting that NrCAM may be more important for inhibitory control at perisomatic sites than at distal dendrites [23]. It was proposed that NrCAM on astrocytes binds neuronal NrCAM at inhibitory postsynaptic sites to mediate synaptic organization, but the interneuron cell type was not identified. These findings prompted us to examine whether NrCAM regulates the density of inhibitory synapses made by CCK- or PV-BCs onto cortical PNs. We focused on the medial prefrontal cortex (mPFC), which includes the anterior cingulate, prelimbic, and infralimbic cortex, because of its essential role in social and cognitive behavior [10–14]. Layer II/III of the mPFC was prioritized, because neurons in this layer are pivotal in integrating long and short range inputs important for such behaviors [24]. Here we demonstrate that NrCAM and AnkB promote the formation, stabilization, or maintenance of perisomatic synapses made by CCK-BCs, but not PV-BCs, onto PNs in the mouse mPFC, implicating NrCAM interactions in development or maintenance of CCK BC to PN connectivity in the mPFC.

## 2. Materials and Methods

### 2.1 Mice

NrCAM-null mutant mice on a mixed 129S6/SvEv^Tac^ x Swiss Webster (CFW) background [25] were maintained by heterozygous matings and bred for multiple generations onto C57Bl/6 mice [18]. WT and NrCAM homozygous-null littermates were used for experiments. Nex1Cre-ERT2: Ank2*^flox^*: EGFP^flox^ (C57Bl/6) conditional mutant mice delete the Ankyrin 2 gene, *Ank2,* from cortical pyramidal neurons under control of endogenous Nex1 regulatory sequences induced by tamoxifen (TMX) [26]. The Nex1 promoter drives TMX-inducible Cre-ERT2 recombinase only in postmitotic PNs and not interneurons, oligodendroglia, astrocytes, or non-neural cells [27]. To generate a reporter mouse line expressing the dsRed variant tdTomato (tdT) in CCK-BCs, Synuclein-gamma (Sncg)-IRES2-FlpO driver mice (JAX 034424; C57Bl/6) [28, 29] were intercrossed with Ai65F mice (frt-stop-frt tdTomato; JAX 032864). Sncg-IRES-FlpO:tdT double heterozygotes for Sncg and tdT alleles were intercrossed to obtain double homozygotes. Offspring were genotyped by PCR and found to be present in Mendelian ratios. Mice were fertile and viable through at least postnatal day P150. All studies used WT and mutant mice balanced for sex. Mice were bred and maintained according to policies of the University of North Carolina Institutional Animal Care and Use Committee (IACUC; AAALAC Institutional Number: #329; ID# 18-073, 21-039) in accordance with NIH guidelines.

### 2.2 Immunofluorescence Staining and Antibodies

Mouse brains were harvested after pericardial perfusion with 4% paraformaldehyde (PFA) in phosphate buffered saline (PBS; NaCl 137 mM, KCl 2.7 mM, Na_2_HPO_4_ 10 mM, KH_2_PO_4_ 1.8 mM). In all experiments 3-5 mice per genotype were used. Coronal sections (40μm) through the PFC were cut with a Leica VT1200S vibratome, and stored at 4°C in PBS with 0.02% sodium azide. Brain sections were placed in a 12-well dish, washed 3×10 min in PBS, then blocked and permeabilized for 1h in a solution of PBS, 10% normal donkey serum (Millipore Sigma 566460) or 5% normal donkey serum with 5% normal goat serum, and 0.3% Triton X-100 (CAS No. 9036-19-5). A second set of 3×10 min washes in PBS followed. The sections then incubated in primary antibody solution for 48h at 4°C. Primary antibody solutions were prepared from in PBS, 0.3% Triton X-100, and 5% normal serum. The negative control well received 1 mL of this solution. Primary antibody solutions contained 1:100 mouse anti-VGLUT3 antibody (Synaptic Systems 135211, 1mg/ml) and 1:100 rabbit anti-Math2/NeuroD6 antibody (Abcam 85824, 0.9mg/ml); 1:500 mouse anti-Parvalbumin (PV) 3C9 antibody (Novus Biologicals NBP2-50038, 1mg/ml) and 1:500 rabbit anti-VGLUT3 antibody (Synaptic Systems 135203, 1mg/ml); or 1:2500 rabbit anti-NECAB1 antibody (Sigma HPA023629, 0.20mg/ml). Mouse monoclonal antibody to vesicular GABA transporter (VGAT) was from Synaptic Systems (131 011.) NrCAM mouse antibody (1:250) was from Biolegend (MMS- 5267) and NrCAM rabbit antibody (1:500) was from Abcam (ab24344). Mouse monoclonal antibodies to synaptotagmin 2b (Syt2b; 1:500) were from the Zebrafish International Resource Center #ANZNP1. Rabbit anti-RFP antibody (Rockland 600-301-379S; 1:1000) and chicken anti-mCherry antibody (Aves Labs MC88697977; 1:250) were used to detect tdTomato.

Brain sections were incubated with primary antibodies at 4°C for 48 hours then washed with PBS 5×10 min. Secondary antibody solutions were prepared in PBS and used at: 1:500 Alexa Fluor 555 donkey anti-mouse antibody (Invitrogen A32773); 1:1000 Alexa Fluor 488 donkey anti-rabbit antibody (Jackson Immunoresearch 711-545-152); 1:667 Alexa Fluor 488 donkey anti-mouse antibody (Invitrogen A32766TR); 1:500 Alexa Fluor 555 donkey anti-rabbit antibody (Invitrogen A31572); 1:1000 Alexa Fluor 647 donkey anti rabbit (Invitrogen 2181018); 1:1000 Alexa Fluor 488 goat anti-chicken (Invitrogen A32931TR). Sections were incubated at 4°C with rocking for 2 hours, then washed 5×10 min with PBS. Sections were removed from wells and placed on Fisherbrand Superfrost Plus microscope slides (Cat. No. 12-550-15, 25 × 75 × 1 mm) with Corning High Precision No.1.5H coverslips (170 ± 5um, 24 × 50mm). Invitrogen Prolong Glass solution (2525808) was used to mount the coverslips. Slides were placed in the dark for 18 to 48 hours, then moved to 4°C.

### 2.3 Imaging Acquisition and Analysis

Images used for quantitative analysis of puncta and soma, and immunolocalization were taken with a Zeiss LSM 700 Confocal Laser Scanning Microscope at the UNC Microscopy Services Laboratory (Pablo Ariel, Director). Identical settings were used to capture images from WT and mutant brain sections. Images were acquired using a pinhole size of 1 AU with pixel sizes of 0.13-0.14 µm. Deconvolution of confocal images was performed utilizing AutoQuant 3 software (Media Cybernetics). For endogenous expression of tdTomato representative images were obtained by wide-field fluorescence microscopy with a 555 nM LED and Semrock filter set for tdT/mCherry. Images were analyzed in FIJI. In all experiments 3-5 mice per genotype were used.

Coronal brain sections through the PFC were subjected to immunofluorescence staining of VGLUT3 or Syt2b, and Math2 or EGFP using Alexafluor-conjugated secondary antibodies. Perisomatic synaptic puncta in the mPFC layer II/III were scored blind to observer on PN soma of WT and NrCAM-null mice, or on EGFP-expressing PN soma in Nex1Cre-ERT2:RCE and Nex1Cre-ERT2: Ank2 F/F: RCE mice. Single optical sections or z-stacks were acquired using confocal microscopy. Image analysis of puncta and soma was conducted using FIJI (ImageJ). Background subtraction was applied to the soma layer using a rolling ball algorithm with a radius of 50 pixels (4.25 μm). The puncta layer was enhanced through Contrast Limited Adaptive Histogram Equalization (CLAHE) (bin size: 24 pixels, 2 μm; 256 bins; maximum slope: 3.00), followed by an unsharp mask filter (radius: 3 pixels, 0.25 μm; mask weight: 0.7) and rolling ball background subtraction (radius: 24 pixels, 2 μm). Puncta were segmented by thresholding the top 4.5% of pixel intensities, and punctate structures were identified using the Analyze Particles function (size range: 0.4–20 μm²; circularity: 0.00–1.00). The detected puncta were overlaid onto the original image, and those within 1.5 μm of the soma boundary were quantified. Puncta were quantified on each image whose magnification and size of field of view are indicated in the Figure legends. Puncta per soma from multiple images were averaged to yield means ± SEM. The figures show representative regions from the original images.

Immunofluorescence labeling of somatic markers NECAB1 and PV was employed to quantify the density of CCK-BCs and PV-BCs, respectively, in layers II/III of the WT and NrCAM-null mPFC. Images for quantitative analysis of soma density were acquired with an Invitrogen EVOS M7000 Imaging System using 10x or 20x objectives. In FIJI, images were processed to minimize signal-to-noise, using rolling ball background subtraction at a 50 pixel radius and median filter at a 10 pixel radius. Soma were identified with the Analyze Particles algorithm (size range: 350–2000 μm²; circularity: 0- 1) and overlaid as outlines onto the original image for counting. NECAB+ soma were counted and densities calculated by dividing the quantity of positive soma by region area in mm^2^. Cell densities per unit area in each image were averaged across multiple images for each genotype. Means were compared by two-tailed t-tests, p < 0.05.

## 3. Results

### 3.1 Perisomatic Synaptic Puncta of CCK-BCs on Pyramidal Neurons are Decreased in NrCAM-null Prefrontal Cortex

NrCAM is expressed in layer II/III PNs in the cingulate region of the mPFC [16] and in interneurons in mouse cortex [23]. A role for NrCAM in the formation of CCK-BC synapses in the perisomatic domain of PNs in the mPFC was evaluated by analyzing CCK-BC synaptic puncta in WT and NrCAM-null mice. CCK-positive, vasoactive intestinal peptide (VIP)-negative basket cells selectively express vesicular glutamate transporter 3 (VGLUT3) at their presynaptic terminals [30], while PNs express the transcription factor Math2/NeuroD6. Immunofluorescence staining of VGLUT3-containing CCK-BC synaptic puncta on Math2+ PN soma showed a significant reduction in the mean number of perisomatic synapses of NrCAM-null PNs compared to WT PNs in adult mice (∼P60) (Fig. 1 A, B; t-test = 0.01). Vesicular GABA transporter (VGAT) is responsible for the uptake and storage of GABA by synaptic vesicles in multiple classes of interneurons. Immunofluorescence staining of VGAT-containing synaptic puncta on Math2+ PN soma showed a smaller but significant reduction in perisomatic synapses in NrCAM-null mPFC compared to WT (Fig. 1 C,D; t-test, p = 0.02).

**Figure 1.**
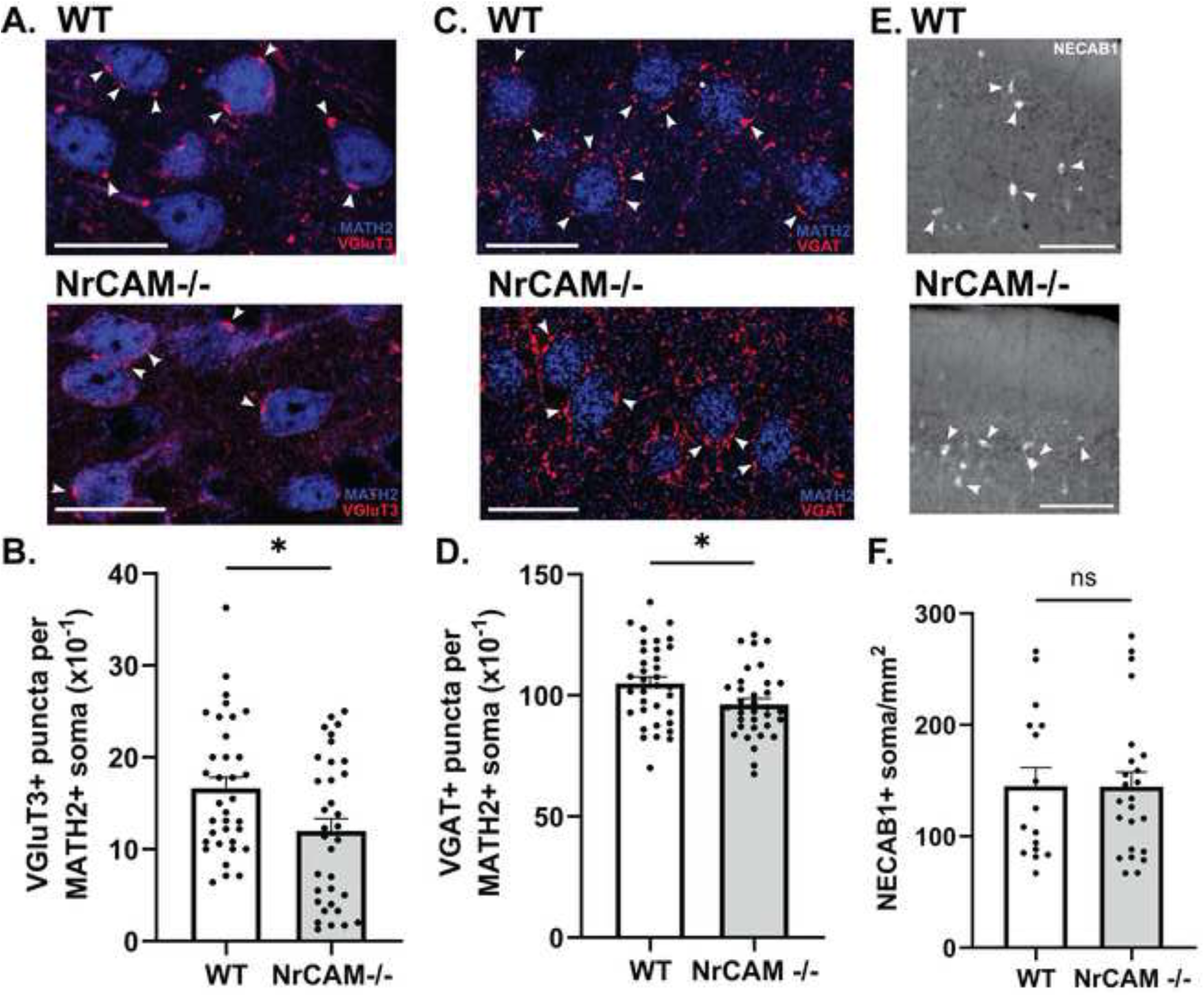
CCK-basket cell synaptic puncta on soma of cortical pyramidal neurons (PNs) are decreased in mPFC of NrCAM-null mutant mice. A. PNs in mPFC layer 2/3 of WT and NrCAM-null mice (adult P60) were immunostained for somal marker MATH2 (blue). CCK-BC presynaptic puncta on PN soma were immunostained for VGLUT3 (red, arrowheads). Confocal images were captured with a 40X objective in a field of view of 1878 x 1878 pixels, or 160 µm x 160 µm. The mean numbers of soma per image were: WT 26 ± 11, NrCAM-null 22 ± 7. Representative regions of confocal images were obtained from maximum intensity projections of 5 optical sections (∼0.3 µm each). Bars = 20 µm. B. Quantification of VGLUT3+ synaptic puncta per MATH2+ PN soma in WT and NrCAM-null mPFC. Data points represent the number of puncta per soma per image. The mean number of puncta per soma across all images was significantly different between genotypes (two-tailed t-test: p* = 0.01). C. PNs were immunostained for MATH2 (blue) and perisomatic puncta from CCK-BCs and PV-BCs were immunostained for VGAT (red, arrowheads) in WT and NrCAM-null mice. Confocal mages were captured with a 63X objective in a field of view of 2048 x 2048 pixels, or 102 µm x 102 µm. The mean numbers of soma per image were: WT 6 ± 2, NrCAM-null 6 ± 2. Representative images as described in A are shown. Bars = 20 µm. D. Quantification of VGAT+ synaptic puncta per MATH2+ PN soma in WT and NrCAM-null mPFC. Data points represent the number of puncta per soma per image. The mean number of puncta per soma were significantly different between genotypes (two-tailed t-test: *p = 0.02). E. Immunostaining of CCK-BC marker NECAB1 (white, arrowheads) in WT and NrCAM-null mPFC. Non-confocal images were obtained by EVOS microscopy. Images were captured with a 10X objective in a field of view of 2048 x 1536 pixels, or 632 µm x 474 µm. The mean numbers of soma per image were: WT 11 ± 4, NrCAM-null 10 ± 4. Representative regions of images are shown. Bar =100 µm. F. Quantification of the mean number of NECAB1+ soma per mm^2^ in EVOS images. Data points represent the number of puncta per soma per image. Differences between WT and NrCAM-null genotypes were not significantly different (ns, two-tailed t-test: p = 0.99).

To control for possible CCK-BC loss caused by NrCAM deletion, which could account for the decreased number of VGLUT3+ perisomatic synapses, antibodies directed against the somal marker N-terminal EF-hand calcium binding protein-1 (NECAB1) were used to immunolabel CCK-BC soma. NECAB1 is located in the nucleoplasm and cytosol of CCK-BCs in cortex and hippocampus, including VGLUT3+/VIP- and VGLUT3+/VIP+ interneurons, a subpopulation of PNs [31], and few PV-expressing interneurons [32]. Quantification of NECAB1-labeled soma showed that there was no significant difference in the mean density of these neurons in layer II/III of WT and NrCAM- null mPFC (Fig. 1 E,F; p=0.99). These results suggested that the reduction of VGLUT3+ perisomatic synaptic puncta on PNs in NrCAM-null mPFC was not due to substantial loss of CCK-BCs.

### 3.2 Perisomatic Synaptic Puncta of Parvalbumin-Expressing Basket Cells are not Decreased in NrCAM-null Prefrontal Cortex

CCK-BCs and PV-BCs are the major interneuron cell types that form perisomatic inhibitory synapses with PNs, and they can target the same PN [4]. Synaptotagmin-2b (Syt2b) is a synaptic vesicle protein involved in calcium-dependent neurotransmission that is a selective marker for PV-BC terminals [33]. To assess whether NrCAM plays a role in perisomatic targeting of PV-BC inputs to PNs, the number of PV-BC synaptic puncta was quantified in the perisomatic compartment of PNs by Syt2b and Math2 double immunofluorescence staining in WT and NrCAM-null mPFC (∼P60). Quantitation of perisomatic puncta in layer II/III revealed no significant difference in the mean density of Syt2b+ synaptic puncta on Math2+ PN soma between the WT and NrCAM-null mPFC (Fig. 2 A,B; p = 0.08). This result indicated that PV-BC synapses onto PN soma were not affected by loss of NrCAM.

**Figure 2.**
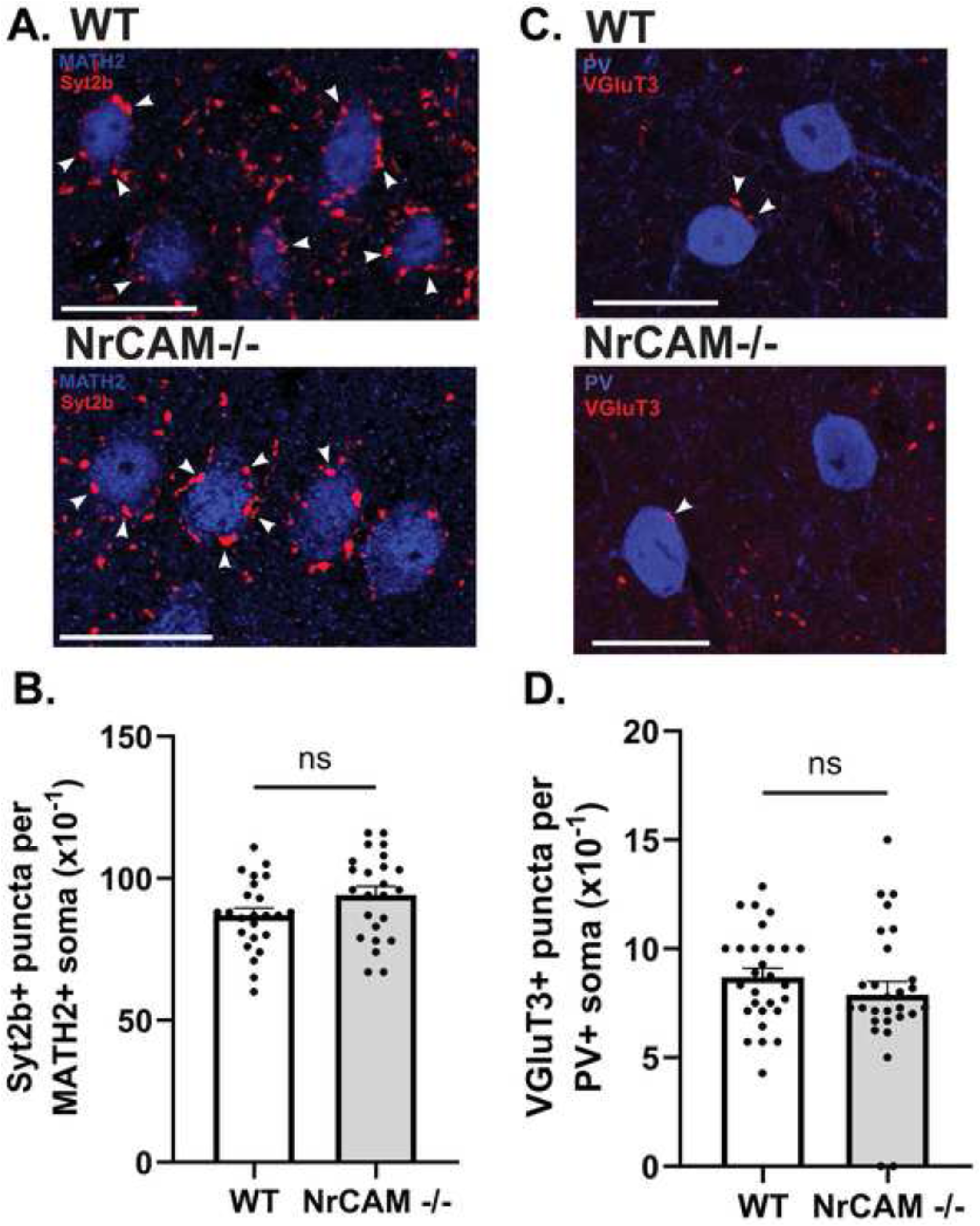
Perisomatic synaptic puncta from PV-BCs onto PN soma are not decreased in mPFC of NrCAM-null mutant mPFC. A. Immunofluorescent staining of Syt2B+ synaptic puncta (red, arrowheads) on MATH2+ PN soma (blue) in WT and NrCAM-null mPFC layer 2/3. Confocal images were captured with a 63X objective in a field of view of 2048 x 2048 pixels, or 102 µm x 102 µm. The mean numbers of soma per image were: WT 10 ± 3, NrCAM-null 10 ± 2. Representative regions of images shown were obtained from single optical sections (∼0.3 µm) (bars = 20 µm). B. Quantitation of Syt2b+ synaptic puncta per MATH2+ soma (x10^-1^). Data points represent the number of puncta per soma in each image. Mean numbers of Syt2b+ puncta per MATH2+ soma between genotypes WT and NrCAM-null were not significant (ns) two tailed t-test: p = 0.08. C. Immunostaining of VGLUT3+ CCK-BC synaptic puncta (red, arrowheads) on PV- BC soma immunostained for parvalbumin (blue). Confocal images were captured with a 40X objective in a field of view of 512 x 512 pixels, or 160 µm x 160 µm. The mean numbers of soma per image were: WT 6 ± 1, NrCAM-null 7 ± 3. Representative regions of images shown were obtained from maximum intensity projections of z-stacks, 5 optical sections (∼0.3 µm each) (bars = 20 µm). D. Quantitation of VGLUT3+ synaptic puncta on PV-BC soma stained for parvalbumin. Data points represent the number of puncta per soma per image. Mean numbers of VGLUT3+ puncta per PV+ soma were not significantly different (ns) between WT and NrCAM-null genotypes (two tailed t-test: p = 0.28).

CCK-BCs target not only PNs but to a lesser extent, the perisomatic compartment of PV-BCs, potentially causing disinhibition [34, 6]. Parvalbumin is predominantly localized to cell bodies of PV-BCs. To determine if NrCAM mediates CCK-BC connectivity with PV-BCs, the effect of NrCAM deletion on the number of VGLUT3+ puncta on PV-BC soma was analyzed. VGLUT3 and PV antibodies were used for double immunofluorescence staining of WT and NrCAM-null mPFC. VGLUT3+ puncta were quantified on PV+ soma in layer II/III. Quantitation of double-labeled puncta revealed no significant difference in the mean density of CCK-BC synaptic puncta on PV-BC soma in WT compared to NrCAM-null mPFC (Fig. 2 C,D; p = 0.28). These results indicated that NrCAM was not significantly involved in the formation or maintenance of CCK-BC inputs to the perisomatic compartment of PV-BCs.

### 3.3 Ankyrin B is Required for CCK-Basket Cell Targeting to Pyramidal Cell Soma

Ankyrins are intracellular scaffold proteins that promote synaptic targeting and stabilization of adhesion molecules, receptors, and ion channels at the neuronal surface [35]. For example the AnkB binding site of L1-CAM is required for targeting of GABAergic inputs to PN soma [36]. Ankyrin G stabilizes somatodendritic GABAergic synapses of cortical and hippocampal interneurons by preventing endocytosis of GABA_A_ receptors [37]. We therefore asked whether AnkB on PNs participated in stabilization or formation of CCK-BC synapses on PN soma. We took advantage of Nex1Cre-ERT2: Ank2*^flox/flox^*: EGFP^flox^ mutant mice, which can be induced to delete *Ank2* specifically from postnatally developing PNs by TMX treatment at P10-P13 [26]. This simultaneously induces EGFP expression and deletes AnkB from PNs, as demonstrated previously by quantitative immunofluorescence and Western Blotting [26]. Nex1-Cre is activated only in postmitotic PNs and not in interneurons or glia [27]. Immunofluorescence staining for VGLUT3 showed a pronounced decrease in synaptic puncta on EGFP+ PN soma in mPFC layer II/III of Nex1Cre-ERT2: Ank2*^flox/flox^*: EGFP^flox^ mice compared to WT Nex1Cre-ERT2: Ank2*^+/+^*: EGFP^flox^ mice (P60-P67) (Fig. 3 A,B; p = 0.001). There was no loss of CCK-BCs in the mutant mPFC, as identified by immunolabeling of NECAB1 (Fig. 3 C,D; p=0.25). Furthermore, the density of Syt2B+ PV-BC puncta on EGFP+ PN soma was unaltered by AnkB deletion from PNs (Fig. 3 E,F; p = 0.99). A large majority (88%) of perisomatic VGLUT3+ puncta on EGFP+ PNs in the mPFC (layer II/III) of Nex1Cre-ERT2: Ank2*^+/+^*: EGFP^flox^ mice co-expressed VGAT, as shown by double labeling of VGLUT3 (magenta) and VGAT (green) (Fig. 3 G). The time of AnkB deletion overlapped with the increase in GAD65+ perisomatic synaptic puncta, which occurs from P10 to P21, and plateaus at P60 in the mPFC [36]. However, the precise timing of axon targeting, synapse formation and synapse stabilization of CCK-BCs is not known. Although cortical interneurons such as PV-BCs derived from the median ganglionic eminence begin to engage PNs in the first postnatal week [38], CCK-BCs which are derived from the caudal ganglionic eminence are delayed relative to PV-BCs [39]. Thus AnkB could play a role in any of these functions.

**Figure 3.**
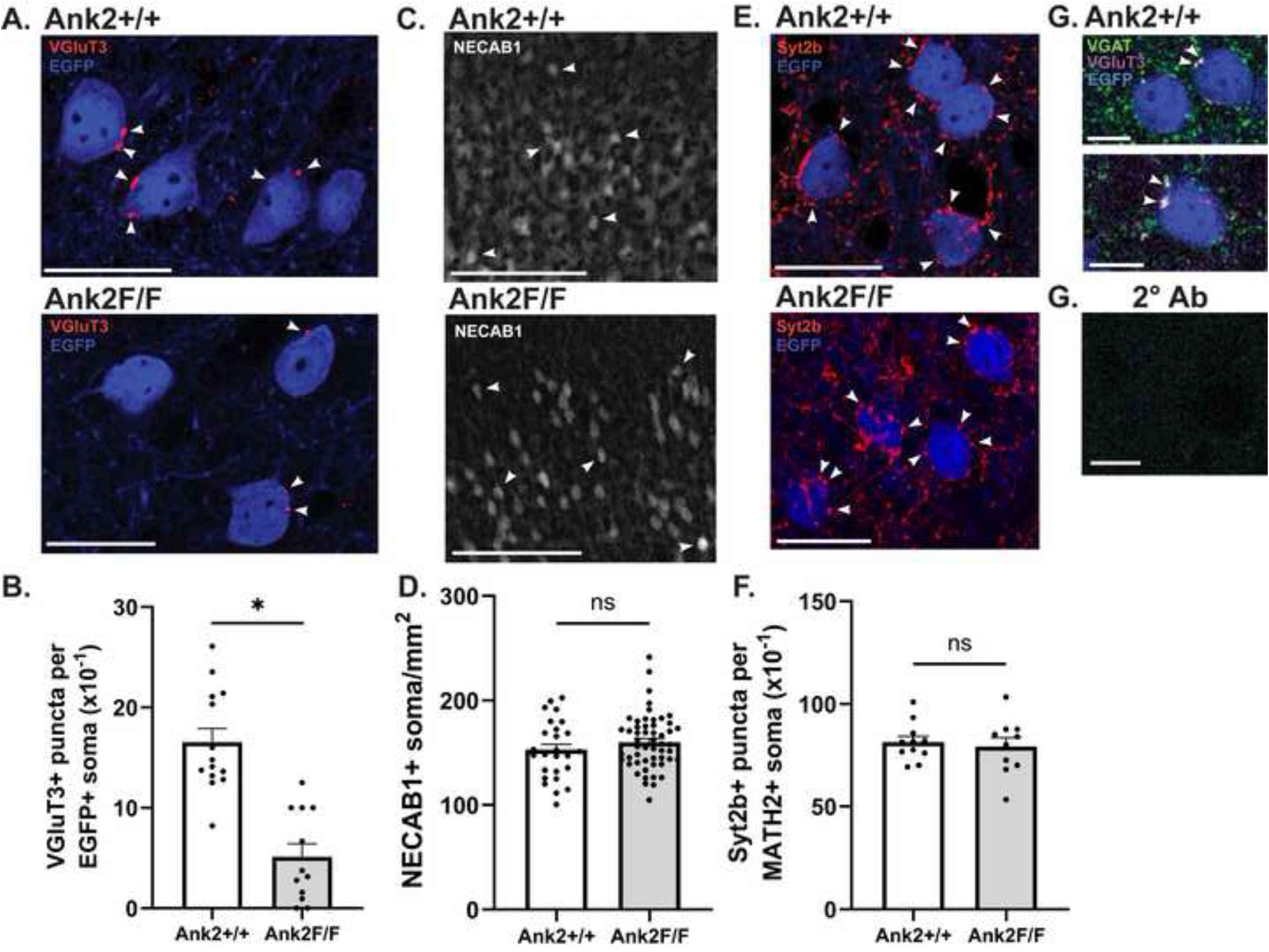
Perisomatic synaptic puncta from CCK-BCs onto PN soma are decreased in mPFC of Nex1Cre-ERT2: Ank2+/+ (WT) and Ank2 F/F mPFC. A. Immunofluorescent staining of VGLUT3+ synaptic puncta (red, arrowheads) on EGFP+ PN soma (blue) in Ank2 +/+ (WT) and Ank2 F/F mPFC layer 2/3 (∼P60). Confocal images were captured with 40X objective and a field of view of 1878 x 1878 pixels, or 160 µm x 160 µm. The mean numbers of soma per image were: WT 18 ± 3, Ank2 F/F 8 ± 1. Representative regions of confocal images were obtained from average intensity projections of 5 optical sections (∼0.3 µm each). Bars = 20 µm. B. Quantitation of VGLUT3+ synaptic puncta on EGFP+ soma. Data points represent the number of puncta per soma for each image. The mean number of VGLUT3+ puncta per EGFP+ soma was significantly different for genotypes Ank2 +/+ and Ank2 F/F (two-tailed t-test, p <0.0001*). C. Immunofluorescent staining of NECAB1+ soma (arrowheads) on EGFP+ PN soma in Ank2+/+ and Ank2 F/F mice. Non-confocal images were obtained by EVOS microscopy. Images were captured with a 10X objective in a field of view of 2048 x 1536 pixels, or 1264 µm x 948 µm. The mean numbers of soma per image were: Ank2 +/+ 12 ± 3, Ank2 F/F 15 ± 4. Representative regions of images are shown. Bars =10 µm. D. Quantitation of NECAB1+ EGFP+ soma/mm^2^ (x10^-1^). Data points represent the number of puncta per soma for each image. Mean differences between Ank2+/+ and Ank2 F/F genotypes were not significant (ns) (two-tailed t-test, p = 0.25.) E. Immunofluorescent staining of Syt2b+ synaptic puncta (red, white arrowheads) on EGFP+ PN soma (blue) in Ank2 +/+ and Ank2 F/F mPFC. Confocal images were captured with a 40X objective in a field of view of 1878 x 1878 pixels, or 160 µm x 160 µm. The mean numbers of soma per image were: Ank2 +/+ 7 ± 3, Ank2 F/F 4 ± 1. Representative regions of images were obtained from maximum intensity projections of 5 optical sections (∼0.3 µm each). Bars = 20 µm. F. Quantitation of Syt2b+ puncta per EGFP+ soma (x10^-1^). Data points represent the number of puncta per soma in each image. Mean differences in Syt2b+ puncta per EGFP+ soma were not significant (ns) between Ank2 +/+ and Ank2 F/F genotypes (two-tailed t-test, p =0.99.) Bars =20 µm. G. Immunofluorescent staining of VGLUT3 (magenta) and VGAT (green) in Nex1Cre-ERT2: Ank2+/+ mPFC layer 2/3 (P63) (EGFP, blue) showed double staining of perisomatic puncta on EGFP+ PNs (white arrowheads), in confocal images, as seen in 2 representative single optical sections (0.3 µm). Quantitation indicated that ∼88% of VGLUT3+ puncta co-expressed VGAT (38 double labeled puncta scored on 43 EGFP+ soma in 8 confocal z-stacks). Control staining with secondary antibodies alone was negative. Confocal images were captured with a 63X objective in a field of view of 512 x 512 pixels, or 160 µm x 160 µm. Bars = 10 µm.

### 3.4 Characterization of a Reporter Mouse Line for CCK-Basket Interneurons

To generate a CCK-BC reporter mouse for localization studies, we took advantage of recent findings that Synuclein-γ (Sncg) is selectively expressed by CCK-BCs but not PV-BCs, somatostatin (SST)- or non-CCK VIP-expressing interneurons [40, 41]. We produced a mouse expressing the fluorescent dsRed reporter tdTomato (tdT) at the Sncg genetic locus. To create this line, Sncg-IRES- FlpO-neo mice (JAX 034434) were intercrossed with Ai65F mice (JAX 032864) containing the Rosa-CAG-Frt-STOP-Frt-tdT sequence. Expression of the recombinase enzyme FlpO in Sncg+ neurons was used to remove the STOP codon and induce tdT expression. Accordingly, brain sections of mice (P60) homozygous for both Sncg-IRES-FlpO and tdT alleles were immunostained for tdT expression using anti-dsRed antibodies. Confocal imaging revealed labeled cells in the mPFC, prominently in layer II/III (Fig. 4 A), as well as in other cortical areas including the primary somatosensory and visual cortex (not shown). Control staining with secondary antibodies alone was negligible (Fig. 4 B). tdT+ cells exhibited a neuronal morphology with labeled soma and emanating processes (Fig. 4 C). Intrinsic tdT fluorescence was much lower but visible in neuronal soma and processes without dsRed antibody enhancement (Fig. 4 D). tdT-expressing cells were also observed in the Sncg-IRES-FlpO:tdT hippocampus and dentate gyrus (not shown). A limitation is that tdT could not be reliably detected in confocal images at perisomatic terminals in the Sncg-IRES- FlpO:tdT mPFC, perhaps due to limited diffusion or transport of tdT in axons.

**Figure 4.**
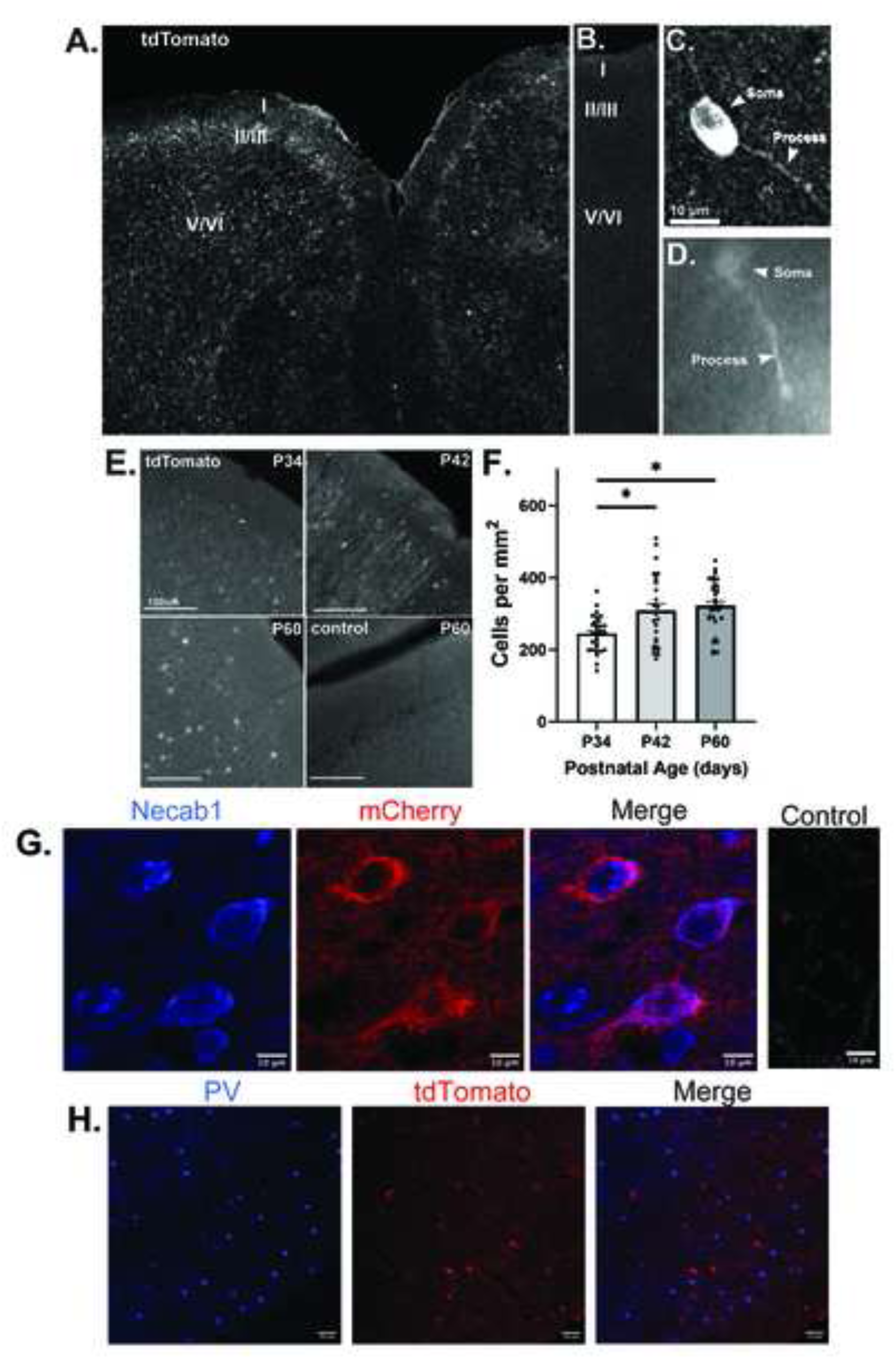
Expression of tdTomato, NECAB1, and Parvalbumin in mPFC of Sncg-IRES- FlpO:tdT mice (P60) A. Immunofluorescent staining of tdT with dsRed antibodies in a representative confocal image (single optical section) captured at 20x. tdT labeled cells were present throughout the mPFC with greater abundance in layer II/III. Cortical layers I-VI are indicated. Image field of view was 1408 x 1408 pixels, or 3904 µm x 3904 µm. B. Control staining with secondary antibodies alone was low. C. Higher magnification of tdT+ cells show neuronal morphology with labeled soma and processes (arrowheads). Bar = 10 µm. D. Intrinsic tdT fluorescence in mPFC layer 2/3 neurons without dsRed antibody staining is lower but visible in soma and processes (arrowheads). E. Time course of tdT expression at postnatal ages P34 (a), P42 (b), and P60 (c, d). Representative nonconfocal images of mPFC layer II/III stained for tdTomato using dsRed antibodies, or secondary antibodies alone, were obtained by EVOS microscopy. Images were captured with a 20X objective in a field of view of 2048 x 1536 pixels, or 633 µm x 475 µm. F. Graph of cell density per mm2 by postnatal age. Significant differences in mean cell density* were determined by Anova and Tukey’s post hoc analysis: P34 vs. P42: p = 0.005*; P34 vs P60: p = 0.0002*; P34-P60: p=0.072, not significant. G. Double immunofluorescence staining for NECAB1 (blue) and tdT using mCherry antibodies (red) in confocal images (single optical sections) showed that a subset of mCherry+ neurons co-expressed NECAB1 (∼25%). Images were captured with a 20X objective in a field of view of 1024 x 1024 pixels, or 319 µm x 319 µm. Representative regions of the confocal images are shown. Bars = 10 µm. Control staining with secondary antibodies alone was negative. H. Double immunofluorescence staining for parvalbumin (PV; blue) and tdTomato (anti-dsRed, red) indicated no detectable co-localization in representative confocal images. Images were captured with a 10X objective at 1024 x 1024 pixels, or 639 µm x 639 µm. Representative regions of confocal images are shown. Bars = 50 µm.

Sncg-IRES-FlpO:tdT mice were also analyzed in a time course for tdT expression by immunofluorescence staining in the developing mPFC (layer II/III) in juvenile (P34-42) and adult (P60) mice (Fig. 4 E,F). Cells expressing tdT showed a significant increase between P34 and P42, and between P34 and P60. The difference between P42 and P60 was negligible, indicating that expression remained fairly constant during later stages of postnatal maturation. tdT+ cells were also present in this region at P21 (Fig. 5). Endogenous Sncg expression in mouse telencephalon/cortex has not been described in detail, but has been shown to arise postnatally, increasing from birth through P49 and adulthood, and is not detectable in embryos (E11.5-E18.5) [42, 43], in general agreement with tdT expression in Sncg-IRES-FlpO:tdT mice.

**Figure 5.**
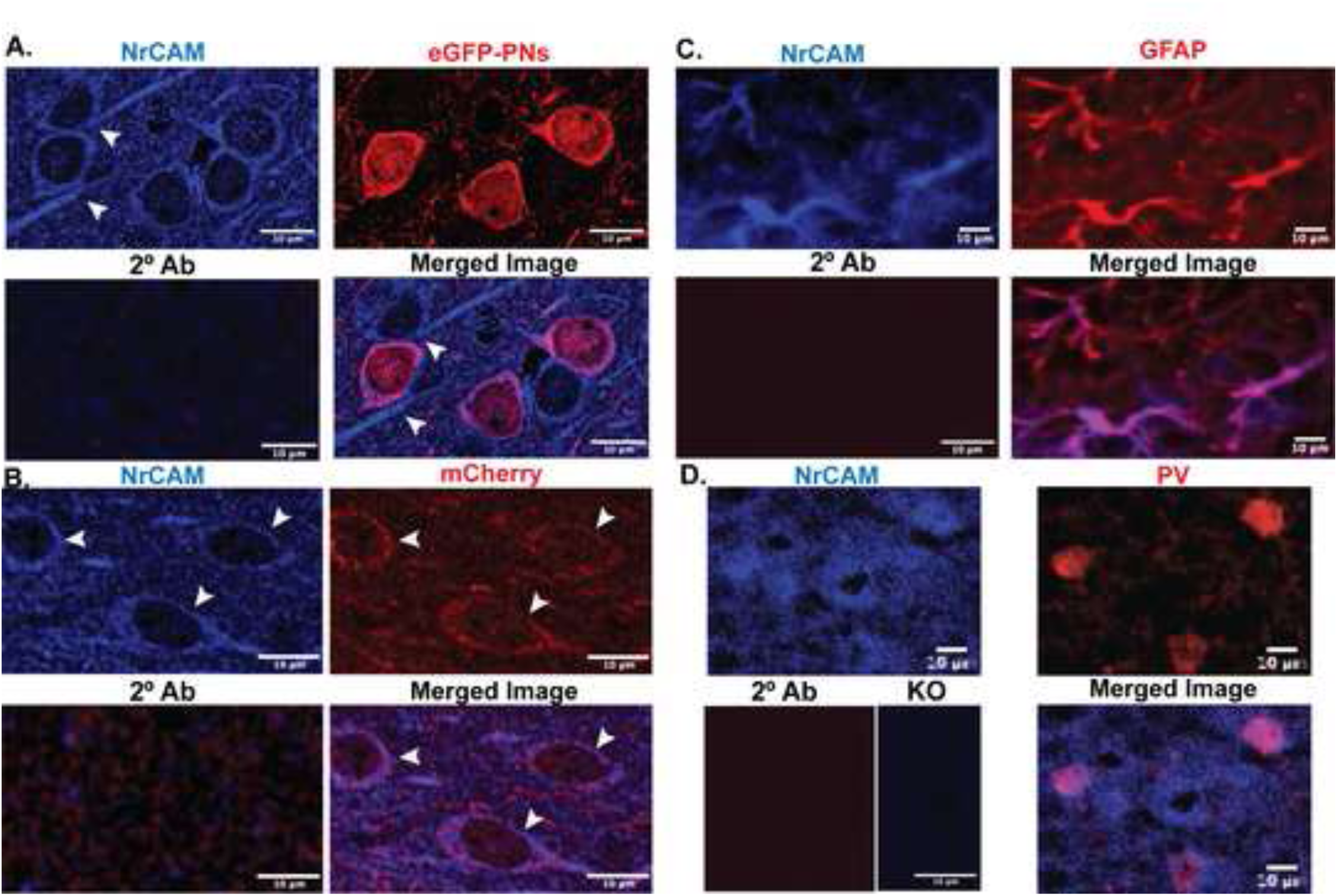
NrCAM expression in pyramidal neurons, CCK basket cells, and GFAP+ astrocytes in mPFC (layer II/III) A. Double Immunofluorescence staining for NrCAM (blue) in Nex1Cre-ERT2: Ank2+/+; EGFP (WT) mice (P21) in PNs expressing eGFP (red). Confocal images captured at 63X (single optical section, 0.3 µm) showed colocalization of NrCAM on PN soma. Arrowheads point to NrCAM+ processes in proximity to EGFP+ PN soma. Control staining with secondary antibodies alone. Bars = 10 µm. B. Double immunofluorescence staining for NrCAM (blue) in tdT+ CCK-BCs of Sncg-IRES-FlpO:tdT mice labeled with mCherry antibodies (red). Confocal images captured at 63X (single optical section, 0.3 µm) show NrCAM localization on CCK-BCs (arrowheads). Control with secondary antibodies alone. Bars = 10 µm. C. Double immunofluorescence staining for NrCAM (blue) and GFAP (red) in tdT+ CCK-BCs of Sncg-IRES-FlpO:tdT mice. EVOS images captured at 20X show colocalization of NrCAM in GFAP+ astrocytes. D. Double immunofluorescence staining for NrCAM (blue) and parvalbumin (PV) (red) in CCK-BCs of Sncg-IRES-FlpO:tdT mice in mPFC layer 2/3. EVOS images captured at 20X show colocalization of NrCAM in a subpopulation of PV+ interneurons. Control staining with secondary antibodies alone was negative. NrCAM antibodies did not label cells in NrCAM-null mice (knockout, KO) as shown in mPFC layer 2/3. Bars = 10 µm.

To further assess reporter expression in CCK-BCs, double immunostaining for tdT with anti-mCherry antibodies (recognizing tdT) and the somal marker NECAB1 was carried out in Sncg-IRES-FlpO:tdT brain sections (P60). Quantitation of double labeled cells in mPFC layer II/III showed that a subset (∼36%) of mCherry+ neurons co-expressed NECAB1 (n=10 confocal images; 39 Sncg+ cells) (Fig. 4 G). mCherry+ cells that did not express NECAB1 may represent a subclass of Sncg+ interneurons, or astrocytes. For example, transcriptomic studies in mouse motor cortex [44] identified 6 distinct subclasses of Sncg+ large and small basket cells, most of which have not been characterized for NECAB1 expression. Scng is also expressed by astrocytes, which do not express NECAB1 [45]. Furthermore, some Sncg+ cells may not express tdT due to incomplete expression of FlpO from the IRES sequence. To further assess the specificity of tdT expression, double immunofluorescence staining was carried out in Sncg-IRES-FlpO:tdT brain sections with anti-PV and anti-dsRed antibodies (Fig. 4 H). Scoring of labeled cells in mPFC layer II/III showed that none of the PV+ neurons detectably expressed dsRed (n=156 cells, 3 mice, 0 double labeled cells).

In summary, characterization of Sncg-IRES-FlpO:tdT cortex indicated that VGLUT3+ CCK-BCs, and not PV-BCs, expressed the tdT reporter.

### 3.5 NrCAM is Expressed by CCK-basket Interneurons and Pyramidal Neurons in the Prefrontal Cortex

To determine if NrCAM localized to PN soma during development when NrCAM expression is highest (∼P21), Nex1Cre-ERT2: Ank2*^+/+^*: EGFP^flox^ mice were induced for EGFP expression (P10-P13) by PNs and immunostained for NrCAM (blue) and EGFP (red). Confocal imaging showed localization of NrCAM on EGFP+ PNs in mPFC layer II/III (Fig. 5 A). Discrete synaptic puncta co-labeled for NrCAM and EGFP were difficult to resolve, although NrCAM+ processes could be seen, some of which were adjacent to EGFP+ PN soma (arrowheads). To ask if NrCAM was also expressed in CCK-BCs, brain sections from Sncg-IRES-FlpO:tdT mice (P21) were immunostained for NrCAM and tdT, then imaged confocally. NrCAM immunoreactivity (blue) was prominent on soma of tdT+ basket cells (red), labeled with mCherry antibodies (Fig. 5 B). However, colocalization of NrCAM with VGlut3 at perisomatic terminals was difficult to observe as NrCAM staining at terminals was not resolvable from the abundant NrCAM staining on PN soma.

NrCAM was also expressed by nearly all GFAP+ astrocytes (94% ± 0.01; n=387 GFAP+ cells) in the mPFC of Sncg-IRES-FlpO:tdT mice (P21) (Fig. 5 C). NCAM localization to astrocytes was previously described in other regions of the mouse neocortex [23]. NrCAM was also expressed by a population of PV-BCs in this region (58% ± 0.01; n=112 PV+ cells), as shown by double labeling with NrCAM and PV antibodies (Fig. 5 D). NrCAM antibodies did not label cells in the NrCAM-null mPFC (Fig. 5 D, KO).

Taken together these results were consistent with cellular roles for NrCAM in CCK-BCs and PN soma in formation, stabilization, or maintenance of perisomatic synapses between CCK-BCs and PNs.

## 4. Discussion

Here we show that NrCAM and its binding partner AnkB are required for optimal synaptic contacts between VGLUT3-expressing CCK-BCs and the perisomatic domain of PNs in the mouse mPFC (layer II/III). A significant decrease was observed in VGLUT3+ CCK-BC synaptic puncta on PN soma in NrCAM-null and AnkB mutant mice compared to WT. This decrease was specific for perisomatic-targeting VGLUT3+ CCK-BCs, because neither NrCAM nor AnkB deletion altered the density of synaptic contacts made by Syt2B+ PV-BCs on PN soma. NrCAM deletion also did not decrease the density of VGLUT3+ synaptic contacts made by CCK-BCs onto PV-BCs, and did not result in interneuron cell loss. Results are consistent with a role for the NrCAM-Ankyrin B interaction in formation, stabilization, or maintenance of connectivity between CCK-BCs and principal excitatory neurons in the mPFC.

Proteomic studies identified NrCAM as a neuronal adhesion molecule necessary for GABAergic inhibitory synapse formation in mouse brain [22, 23]. However these reports did not determine the interneuron cell type or subcellular compartment involved. Results shown here support the interpretation that NrCAM promotes synaptic connectivity of VGLUT3+ CCK-BCs, and not PV-BCs, at the perisomatic domain of PNs. The present work demonstrates expression of NrCAM on postnatally developing PNs, CCK-BCs, and astrocytes. Exploiting a new transgenic mouse line Sncg-IRES-FlpO:tdT, which labels a population of VGLUT3-expressing CCK-BCs in the mPFC, we show that NrCAM is expressed by Sncg+ CCK-BCs. In the conditional mouse mutant Nex1Cre-ERT2: Ank2*^+/+^*: EGFP^flox^, NrCAM localized to PN soma and processes. These results are consistent with a role for NrCAM-AnkB in CCK-BC synaptic perisomatic connectivity on PNs. This role accords with the ability of NrCAM to associate with Gephyrin, an inhibitory postsynaptic scaffold protein [22], and with localization of AnkB to the PN somatodendritic compartment [19].

Homophilic or heterophilic interactions of NrCAM with itself or F3/contactin, TAG1, or Neurofascin [46] could mediate adhesive contacts between CCK-BC axons and PN soma, or astrocytes. A limitation is that NrCAM deletion in global null mutant mice affects expression in CCK-BCs, PNs, and astrocytes, thus effects on CCK-BC synapses cannot be ascribed specifically to pre- or postsynaptic neurons, or glia. However, NrCAM in neurons clearly contributes to GABAergic synaptogenesis since Takano et al., [23] showed that NrCAM knockdown in neurons under control of the neuronal promoter Synapsin I decreases the number of inhibitory synaptic contacts. In regard to glia, tripartite neuronal-astrocyte interactions have been implicated in interneuron-PN synaptic connectivity [23]. It is therefore reasonable to speculate that homophilic or heterophilic NrCAM binding between CCK-BCs and PNs, and/or between neurons and glia, helps establish or maintain synaptic adhesion.

NrCAM and its interaction with AnkB may cooperate with other adhesion mechanisms to promote CCK-BC synaptic development. For example, the Ankyrin binding site on L1 is required for stabilizing GABAergic inputs to PN soma in the mPFC [36]. L1 also drives chandelier interneuron innervation of the PN axon initial segment through Ankyrin G anchoring [47]. The synaptic molecule Dystroglycan also participates in perisomatic targeting of CCK-BCs [48]. Finally, Sncg-IRES-FlpO:tdT mice developed here will help advance our knowledge of CCK-BCs, which has been limited by differences in expression in other lines that target CCK-BCs, including CCK-Cre [49, 7], VGLUT3-Cre [8], and cannabinoid-1 receptor Cre [50] mice.

## 5. Conclusion

The findings of this study demonstrate a role for NrCAM and its binding partner AnkB in establishing or maintaining connectivity between VGLUT3- expressing CCK-BCs and PN soma the mouse mPFC. Deletion of NrCAM in null mutant mice resulted in decreased numbers of VGLUT3+ CCK-BC synaptic contacts in mPFC layer II/III, which likely impacts critical cortical networks. Such effects may lead to behavioral deficits in sociability, reversal learning, and addiction vulnerability, all of which have been described in NrCAM mutant mice [21, 51]. The unique property of CCK-BCs in mediating excitatory or inhibitory transmission [8] suggests that functionality of mPFC circuits could be differentially affected by NrCAM loss under high or low glutamatergic tone. It has been postulated that cortical microcircuits in ASD are associated with an increased ratio of excitatory/inhibitory synapses that results in hyperexcitability [52, 53]. Deficiencies in NrCAM and the high confidence ASD risk factor AnkB may contribute to excitatory/inhibitory imbalance by diminishing connectivity between CCK-BCs and principal excitatory neurons in the mPFC, affecting cognitive and social behaviors relevant to ASD.

## Abbreviations

CCK-BCs: cholecystokinin-expressing basket cells
PNs: pyramidal neurons
Sncg: Synuclein-γ
PV: Parvalbumin
VGLUT3: vesicular glutamate transporter 3
VGAT: vesicular GABA transporter
Syt2: synaptotagmin-2
tdT: tdTomato.

## Acknowledgements

We thank Erin Zhang and Ernest Pereira for expert assistance with the experiments. Dr. Paul Manis is gratefully acknowledged for microscopy of intrinsic fluorescence in the reporter mouse line. Dr. Pablo Ariel, Director of the UNC Microscopy Services Laboratory (MSL) in the Departments of Pathology and Laboratory Medicine, provided advice and assistance with confocal microscopy. Dr. Bryce Duncan (UNC) kindly provided help with EVOS microscopy.

